# The central motor command, but not the muscle afferent feedback, is necessary to generate the perception of effort

**DOI:** 10.64898/2026.02.04.703832

**Authors:** Benjamin Pageaux, Maxime Bergevin, Luca Angius, Thomas Mangin, Romuald Lepers, Samuele M. Marcora

**Author notes:** Equally contributed to this work (Benjamin Pageaux, Maxime Bergevin). Corresponding author: Benjamin Pageaux, École de kinésiologie et des sciences de l’activité physique (EKSAP), Centre d’éducation physique et des sports (CEPSUM), Faculté de médecine, Université de Montréal, 2100, boul. Édouard-Montpetit Montréal, QC H3T 1J4 Canada.

## Abstract

Two theoretical models are proposed on the signal processed by the brain to generate the perception of effort (PE): the corollary discharge model and the afferent feedback model. To test the validity of these models, we used electromyostimulation to manipulate the magnitude of the central motor command during *voluntary* (high motor command), *evoked* (no motor command) and *combined* (low motor command) contractions at similar torque outputs. As electromyostimulation evokes sensory volleys to the central nervous system, it was used to evoke muscle contractions and to stimulate afferent feedback. We hypothesized that PE would reflect the magnitude of the central motor command and that evoked muscle contractions in the absence of central motor command would not elicit any PE. Twenty participants (n = 10 experienced, n = 10 novice with electromyostimulation) volunteered in this study. Participants reported their PE after isometric (10% and 20% MVC) and dynamic (5% and 20% MVC) voluntary, evoked, and combined contractions. For the same torque, participants reported no PE during evoked contractions, but all reported PE during voluntary contractions. Experienced but not novice participants reported lower PE during the combined than during voluntary contractions. This study provides evidence that the presence of central motor command is necessary to generate the perception of effort. However, results from the novice participants during the combined contractions suggest that other factors such as inhibitory control may affect PE. Future studies should investigate the relationship between the central motor command and PE during physical tasks at various levels of complexity.

## Introduction

The perception of effort is a common experience in daily life and plays a critical role in regulating human behaviour (Preston & Wegner, 2009). Perception of effort can be described as “*the particular feeling of that energy being exerted, or the phenomenal experience of effort, […] a sensation of strain and labor, a feeling that intensifies the harder a person tries.*” (Preston & Wegner, 2009). In the context of physical exercise, effort perception has been defined as the *conscious sensation of how heavy, hard and strenuous a physical task is* (Marcora, 2010). From a motivational science perspective, effort perception acts as a signal that allows both healthy (Grummt et al., 2024) and symptomatic individuals (Brinkmann & Gendolla, 2008) to gauge their engagement in a task. This perception is fundamental to the initiation and regulation of physical activity (Marcora, 2016; Marcora & Staiano, 2010), and shapes human behavior by informing the decision to persist or disengage in a task. Despite its importance in the regulation of human behavior and performance, the neurophysiological mechanisms underlying effort perception remain poorly understood. In particular, the nature of the signal processed by the brain to generate the perception of effort has been extensively debated. The central question is whether this signal originates within the central nervous system or from the periphery. In the cognitive psychology and neuroscience literatures, effort perception is typically attributed to internal signals related to cognitive control and other executive functions (Shenhav et al., 2017). In contrast, in the field of sports science, the debate remains active (Marcora, 2009; Pageaux, 2016), with some arguing that effort perception arises from feedback originating in working muscles (Broxterman et al., 2018), while others propose that it reflects the magnitude of the central motor command (De Morree et al., 2012).

To date, two competing models have been used to explain effort perception during physical exercise (Marcora, 2009; Pageaux, 2016). The *afferent feedback model* posits that the brain interprets sensory feedback from the working muscles. This feedback can arise from either group I-II or group III-IV muscle afferents. Group I-II afferents are innervated by specialized mechanoreceptors located in muscle spindles and Golgi tendon organs, and contribute to proprioceptive feedback, motor control, and the perception of position, movement, force and muscle tension (Proske & Gandevia, 2012). Their role in effort perception remains unclear, although some evidence suggests that selectively impairing Ia afferent feedback reduces perceived effort during elbow flexion (Monjo et al., 2018) and cycling (Marchand et al., 2025). However, because γ-motoneuron couples spindle activity to the motor command (Dimitriou, 2022), it is inherently difficult to determine whether group Ia afferents generate or modulate perceived effort during voluntary contraction (Monjo & Allen, 2023). Group III-IV afferents are innervated by free nerve endings that contribute to cardiorespiratory regulation (Amann et al., 2020) and transmit nociceptive signals (Pollak et al., 2014). Several studies have shown that attenuating feedback from group III-IV afferents reduces ratings of perceived effort during knee extensions (Broxterman et al., 2018) and cycling (Amann et al., 2010), but this effect is typically observed only at high intensity (Bergevin et al., 2023). At high intensities, metabolite accumulation strongly activates these nociceptive afferents, leading to naturally occurring muscle pain (O’Connor & Cook, 1999; O’Malley et al., 2024). Therefore, these findings may reflect reductions in pain rather than in effort perception, as ratings of perceived effort instructions often included discomfort and pain (Bergevin et al., 2023). Although group I–IV muscle afferents are typically activated during voluntary contractions, their activation can occur independently of the motor command, arising instead from mechanical alterations within the muscle. For instance, on resting muscles, tendon or muscle-belly vibration can elicit illusions of movement and position, and intramuscular injections of metabolites associated with muscle contractions can induce sensations related to fatigue and pain (Pollak et al., 2014). To the best of our knowledge, it remains unknown whether simultaneous stimulation of group I-IV muscle afferents in the absence of motor command can generate the perception of effort.

In contrast, the corollary discharge model proposes that the perception of effort is generated by the brain’s processing of internal copies of motor commands, known as corollary discharges or efferent copies. These discharges originate in motor and pre-motor areas and are sent to sensory regions. They are involved in various processes, including motor control, proprioception, and the modulation of other perceptions (Monjo & Allen, 2023; Proske & Gandevia, 2012). The corollary discharge model does not exclude a peripheral contribution to effort perception, but suggests that afferent feedback indirectly affects, rather than generates it (Bergevin et al., 2023, p. 416; De Morree et al., 2012, 2014). In this framework, the perception of effort reflects the magnitude of the central motor command rather than peripheral sensory feedback. Supporting this view, the motor-related cortical potential, an electroencephalogram index of central motor drive, rises in parallel with ratings of perceived effort during both actual (De Morree et al., 2012) and imagined contractions(Jacquet et al., 2021). Manipulating the motor command experimentally produces corresponding changes in perceived effort. For instance, experimentally induced muscle fatigue increases the required motor command and increases perceived effort (De Morree et al., 2012). Conversely, caffeine ingestion decreases the required motor command and decreases perceived effort (De Morree et al., 2014). Surface electromyography provides another index of the central motor command. When muscles were transiently weakened by prior shortening, participants produced larger electromyographic amplitudes and reported higher effort perception compared to a condition with prior lengthening-induced strengthening (Kozlowski et al., 2021). Notably, when participants were instructed to match electromyographic amplitude rather than torque, perceived effort no longer differed (Kozlowski et al., 2021). Furthermore, during eccentric cycling, both low (30 rpm) and high cadence (90 rpm) elicited higher electromyographic amplitudes and greater perceived effort than a moderate cadence (60 rpm; Mater et al., 2023). Together, these findings consistently show that perceived effort scales with indices of the central motor command.

Electromyostimulation is a technique that involves non-invasive stimulation of the muscle belly via electricity delivered through electrodes placed over the muscle or the motor nerve, thereby inducing muscle contraction through activation of motor axons (Bergquist et al., 2011). This technique can be used to evoke muscle contractions, without engaging central motor command. Electromyostimulation also stimulates sensory fibres and sends a sensory volley to the central nervous system (Bergquist et al., 2011). In this context, group I-II sensory fibres, related to muscle spindles, play a role in proprioception, such as muscle tension (Proske & Gandevia, 2012), whereas group III-IV sensory fibres are associated with the experience of discomfort and pain (Pollak et al., 2014). Therefore, in addition to their regulatory role in motor control during exercise, these fibres reach both subcortical and cortical areas where various perceptions, such as muscle tension and pain, are ultimately generated (Craig, 2003; Proske & Gandevia, 2012). Electromyostimulation can also be used to create combined contractions by superimposing voluntary contractions onto evoked contractions (Descollonges et al., 2025). In the functional electrical stimulation literature, such combined contractions are used to improve movement and reduce the central motor command required to produce a given target torque or power output (Descollonges et al., 2025). At the same time, the afferent feedback is increased due to the direct electrical stimulation of sensory fibres. This allows dissociating the contributions of the motor command and afferent feedback at the same torque level. Importantly, electromyostimulation can be applied during both isometric (Boerio et al., 2005) and dynamic contractions (Descollonges et al., 2025). This distinction is crucial for capturing a broader range of muscle spindle feedback because the human nervous system employs γ-motoneurons during isometric and dynamic contractions (γ-static and γ-dynamic, respectively) to tune spindle sensitivity (Dimitriou, 2022; Kakuda & Nagaoka, 1998). Thus, investigating the perception of effort during isometric and dynamic contractions ensures that afferent feedback specific to muscle length is also considered.

This study aimed to test the validity of the corollary discharge and afferent feedback models. We used electromyostimulation to manipulate the magnitude of the central motor command and afferent feedback during isometric and dynamic contractions at the same torque produced. First, we adopted a deductive approach (Briand et al., 2025) to test the primary hypothesis that the generation of effort perception is dependent on the presence of a central motor command. Therefore, no perception of effort was expected during evoked contractions, in which no central motor command is generated, and high afferent feedback is induced by the electromyostimulation (Bergquist et al., 2011). Second, we explored the relationship between the magnitudes of the motor command, afferent feedback and effort perception by comparing voluntary and combined contractions. We hypothesized that, when a central motor command is present, the perception of effort reflects its magnitude. Accordingly, lower ratings of perceived effort were expected during combined contractions (low central motor command; high afferent feedback) compared with voluntary contractions (high central motor command; low afferent feedback). Healthy individuals show increased tolerance to electromyostimulation with repeated exposure, with pain being a leading cause for intolerance (Alon & Smith, 2005). This is consistent with research showing attenuated pain responses to noxious stimuli following repeated exposure (Bingel et al., 2007). Consequently, individuals who have had prior experiences with electromyostimulation should experience less intense pain when stimulated. Because effort spans cognitive and motor domains, and pain mobilizes attentional resources, making the task more costly (Torta et al., 2017), we included participants who were either novice or experienced with electromyostimulation to explore potential pain-related confounds in effort ratings (Bergevin et al., 2023)

## 2. Methods

### 2.1 Participants

Twenty healthy adults volunteered to participate in this study (mean ± SD; age: 26 ± 9 years, height: 175 ± 10 cm, weight: 69 ± 15 kg, 15 males and 5 females). A sample size of 20 participants was chosen i) to be consistent with similar psychophysical studies in the field of effort perception (De Morree et al., 2012; Kozlowski et al., 2021) and ii) with our possibility to recruit participants in the time allocated to data collection (Lakens, 2022). A detailed sensitivity analysis, related to our a priori hypothesis and exploratory analysis, is available in the section “2.5.1. Sensitivity analyses” below. Half of the participants were already experienced with electromyostimulation, and the other half were novices. An experienced participant was defined as someone who had participated in at least 2 studies involving electrical stimulation or had previously participated in training or rehabilitation involving electromyostimulation. As this information was collected at the beginning of the first visit, the experimenters were not blinded to group assignment during data collection.

Each participant gave written informed consent before the study. Experimental protocol and procedures were approved by the local Ethics Committee of the School of Sport and Exercise Sciences, University of Kent (Prop97_2013_14). The study conformed to the standards set by the World Medical Association and the Declaration of Helsinki, except for preregistration. All participants were given written instructions describing all procedures related to the study, but were naive to its aims and hypotheses. At the end of the last session, participants were debriefed and asked not to discuss the study’s true aim with other participants.

### 2.2 Overview

Participants visited the laboratory on two separate occasions. They were instructed to abstain from alcohol and vigorous activity for 24 hours before each visit, and to avoid caffeine for at least 3 hours before testing. Participants were also asked to sleep at least seven hours the night before each session. The first visit served as a familiarisation session during which participants were introduced to all experimental procedures. During the second visit, participants performed both isometric and dynamic knee extension contractions under three conditions: voluntary, evoked and combined. Participants were randomized to start with either isometric or dynamic contractions. Within each contraction modality, the order of the three conditions (voluntary, evoked, combined) was randomized. Each session began with a standardized warm-up, consisting of 10 brief submaximal contractions at approximately 50% of the participant’s estimated peak torque. Maximal voluntary contraction (MVC) torque was then assessed using two isometric contractions lasting three seconds each. A third MVC trial was performed if the difference between the first two attempts exceeded 5%. Visual feedback showing the torque produced during the previous MVC and verbal encouragement were provided to promote maximal effort. **Figure 1** presents an overview of the protocol.

**Figure 1.**
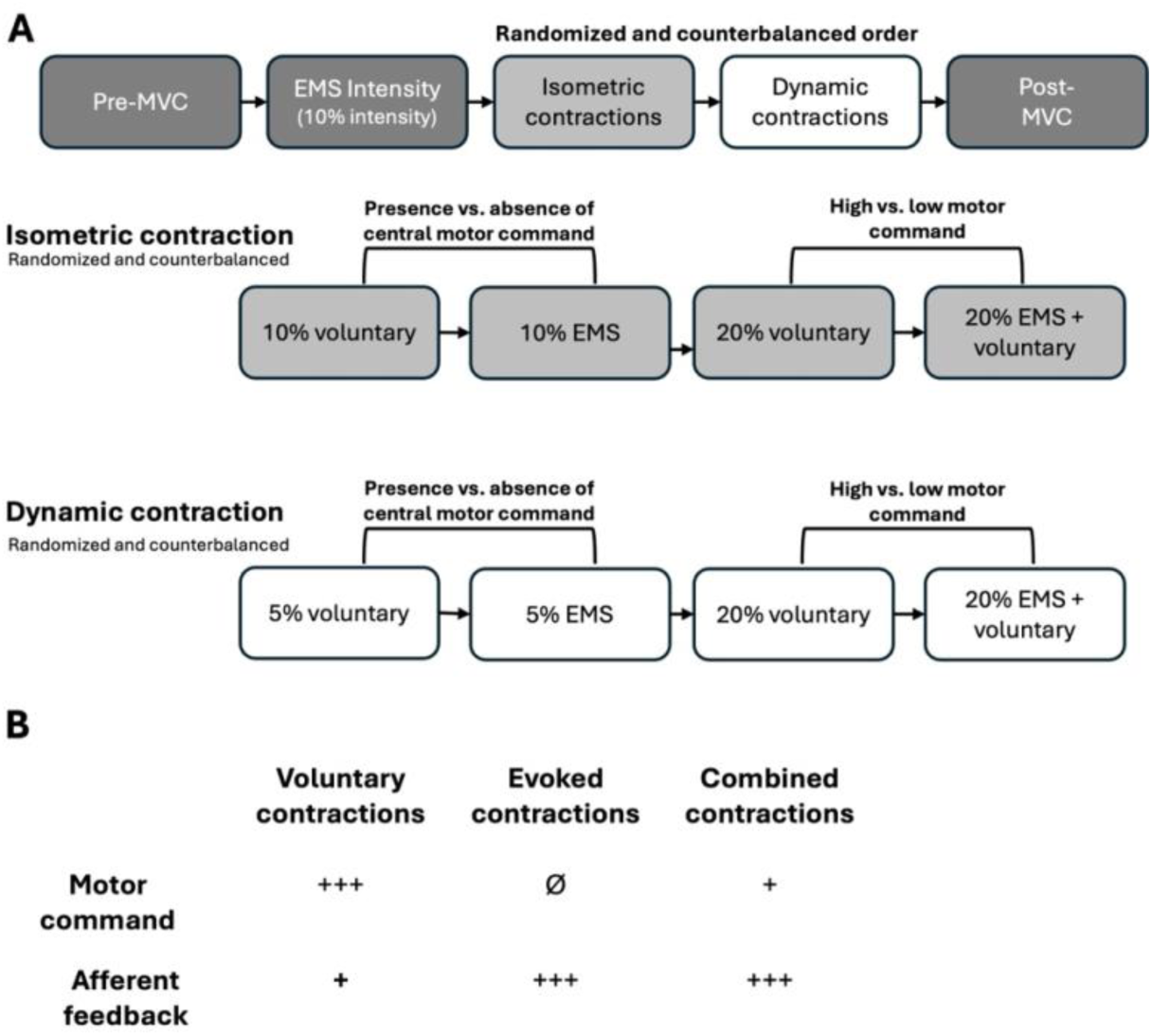
Overview of the protocol. Isometric and dynamic contractions were executed in a randomized order; voluntary, evoked, and combined contractions were also performed in a randomized order (panel A). Representative single-trial traces are displayed alongside each condition to illustrate the general form of each contraction type. For voluntary and evoked contractions, contractions began after the “Go!”. During the combined contractions, the stimulator was set to “1”, and participants were asked to superimpose a voluntary contraction at “Go!” (panel B). Voluntary contractions were characterized by relatively high motor commands and low afferent feedback. Evoked contractions were characterized by the absence of motor command and high afferent feedback. Combined contractions were characterized by relatively low motor commands and high afferent feedback (panel C).

Following MVC assessment, the stimulation intensity required to elicit 10% of the participant’s MVC peak torque was determined (see *Electromyostimulation*). All trials were performed with the right leg. All isometric and dynamic contraction trials were performed separately. To avoid metabolite accumulation, each contraction was separated by a rest period of at least 30 seconds. Participants completed three trials per condition. After each trial, perceptual ratings of effort, force and muscle pain in the working leg were collected during the resting period between trials.

### 2.3 Task

Each trial began with a verbal countdown from the experimenter (“3, 2, 1, GO”). For voluntary contractions, participants were asked to reach the required torque at the “GO” signal. For evoked contractions, participants were instructed to remain fully relaxed as the stimulation was applied at “GO”. During combined contractions, stimulation began at “1”, and participants were instructed to begin applying their portion of the required torque and reach the target torque at “GO”. Participants were not informed of the specific target torques for each contraction.

To isolate the role of the central motor command in effort perception, we compared voluntary contractions to evoked contractions via electromyostimulation. This allowed us to differentiate between contractions driven by the central motor command and those involving afferent feedback. To assess the effect of central motor command magnitude, we compared voluntary contractions to combined contractions. During combined contractions, less motor command is required of the participant, while afferent feedback is increased by the stimulation (Bergquist et al., 2011).

#### 2.3.1 Isometric contractions

Participants performed 5-second isometric knee extensions with the knee fixed at 90° of flexion. For voluntary contractions vs. evoked contractions (i.e., motor command present vs. absent), the target torque was set at 10% of the participant’s MVC. For voluntary vs. combined contractions (i.e., high vs. low motor command), stimulation began at “1” to generate 10% of MVC torque, and participants were instructed to superimpose a voluntary contraction at “GO” to match 20% MVC. In contrast, voluntary contractions required the full 20% MVC to be generated volitionally, involving a greater motor command and less afferent feedback.

A target torque line displayed on a computer monitor positioned approximately 1.5 meters in front of the participants had to be matched. To avoid any bias, participants were blind to the y-axis (torque) values, which were adjusted between trials to keep the target line at the same on-screen position.

#### 2.3.2 Dynamic contractions

Pilot testing revealed that evoked contractions involving a range of motion greater than 40° caused considerable muscle pain. To improve participant comfort, the range of motion was limited to 90°-40°. Participants performed dynamic knee extensions from 90° to 40° of knee flexion (0° = full knee extension). During voluntary contractions, they were instructed to move the dynamometer arm at a constant rate through the full range of motion, applying just enough torque to initiate and maintain the movement.

For voluntary contraction vs. evoked contractions (i.e., high vs. low motor command), the target torque that allowed the dynamometer arm to move was set at 5% of MVC. Stimulation intensity was calibrated to produce 10% MVC at 90°, consistent with the isometric evoked contractions. However, because torque produced by a fixed stimulation intensity declines at the knee extends due to the muscle’s force-length relationship, pilot testing revealed that a 10% MVC torque prevented the dynamometer arm from completing the full range of motion. The movement threshold was therefore set to 5% MVC.

For voluntary contractions vs. combined contractions (i.e., high vs. low motor command), stimulation began at “1” to generate 10% of MVC torque, and participants were instructed to superimpose a voluntary contraction at the “GO” signal to reach a total of 20% MVC. In contrast, voluntary contractions required the full 20% MVC to be generated voluntarily, involving a greater motor command and less afferent input. During these conditions, the target torque that allowed the dynamometer arm to move was set to 20% of MVC. The trial initiation procedure was identical to the isometric task.

**Figure 2.**
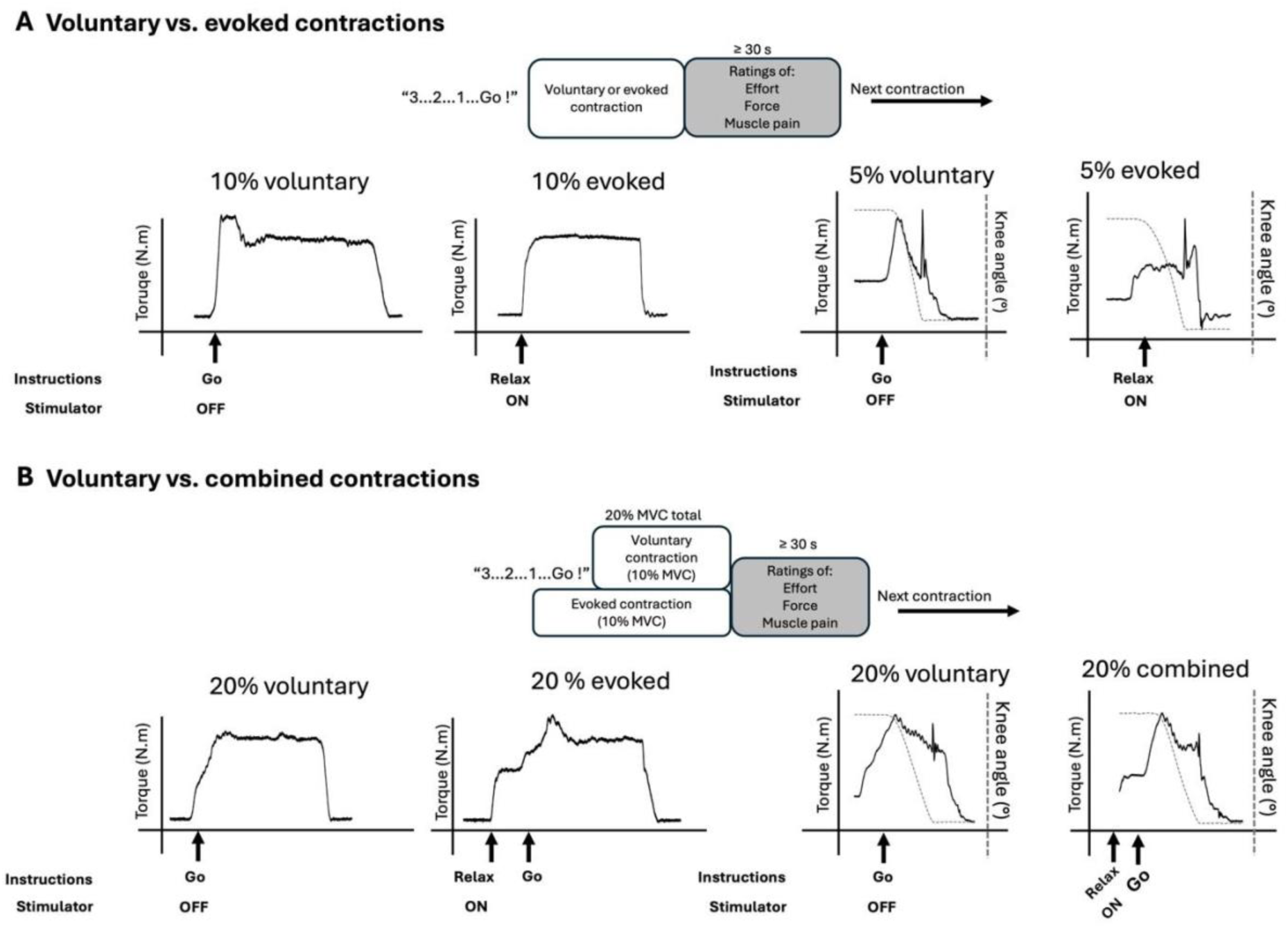
Sequence of events and representative torque traces for voluntary vs. evoked contractions (panel A). The left panel depicts 10% MVC isometric contractions; the right panel depicts 5% MVC dynamic contractions (dashed line indicates knee angle; torque spike represents the limb reaching the mechanical end-stop of the dynamometer). Voluntary contractions (stimulator OFF) were initiated by a “GO” command. Evoked contractions (stimulator ON” were induced while participants remained relaxed. Sequence of events and representative torque traces for 20% MVC voluntary vs. combined contractions (panel B). Participants performed either purely voluntary or combined contractions. During the combined contractions, a 10% MVC evoked contraction was initiated first (stimulator ON), followed by a superimposed 10% MVC voluntary contraction (“GO” command) to achieve the 20% target torque.

#### 2.3.3 Electromyostimulation

A high-voltage constant-current stimulator (maximal voltage 400 V, model DS7 modified, Digitimer, Hertfordshire, UK) was used to apply electromyostimulation on the knee extensor muscles. The knee extensor muscles were stimulated using a pair of surface electrodes (10×5 cm, Phoenix Healthcare Products Ltd., Nottingham, UK) positioned perpendicular to the long axis of the femur. Proximal and distal electrodes were respectively positioned ∼5 cm below the inguinal ligament and ∼10cm above the patella. After pilot testing, a stimulation frequency of 30 Hz was chosen to obtain a torque plateau during the stimulation, allowing the participant to easily superimpose a voluntary contraction in addition to the evoked contraction. The stimulation intensity required to elicit 10% MVC was determined using an iterative procedure. Current intensity was gradually increased until target torque was reached. Following a 30-s rest period, the contraction was repeated to confirm that the same intensity still produced the target torque. Current intensity was adjusted, and this verification step was repeated as needed until the target torque was consistently matched. The intensity producing 10% of the pre MVC was kept constant through the protocol (72.7 ± 12.5 mA).

#### 2.3.4 Mechanical recordings

Torque was recorded using a dynamometer (Cybex ORM isokinetic dynamometer, CMSi, Computer Sports Medicine Inc., Stoughton, USA). During the tests, a two-shoulder harness and a belt across the abdomen limited extraneous movements of the upper body. MVCs and isometric contractions were performed at a knee joint angle of 90° of flexion and a hip angle of 90°. The torque-time integral was calculated offline using AcqKnowledge 4.1 (Biopac Systems Inc, USA) for all contractions as the area under the curve to quantify the intensity-duration and total mechanical work performed.

### 2.4 Perceptual measures

To minimize the risk of perceptual conflation (Gruet et al., 2018), we used three distinct scales for the perceptions of effort, pain and force. The perceived effort intensity was measured using the RPE 6-20 scale (Borg, 1998). As the inclusion of other-exercise sensations (e.g., pain) may bias ratings of perceived effort (Bergevin et al., 2023), participants were also instructed to rate their perceptions of force and pain separately (Pageaux, 2016). Leg muscle pain, defined as “the intensity of hurt that a participant feels in his quadriceps muscles, was measured using the Cook Pain Intensity Scale (O’connor & Cook, 2001). Participants were asked to verbally report their estimation of the force produced using a visual analogue scale (0 = “no force at all” and 100 = “maximal force”). Participants were instructed that the MVC performed at the beginning of the experimental session corresponded to their maximal force, i.e., a rating of 100. All perceptual measurements were taken during the resting period between each contraction.

#### 2.4.1 Perception of effort

Perception of effort, defined as “the conscious sensation of how hard, heavy, and strenuous exercise is” (Marcora, 2010), was measured during the resting period between each contraction using the RPE6-20 scale (Borg, 1998). Standardized instructions for the scale were given to each subject before the warm-up: *“During this experiment, we want you to rate your perception of effort, defined as the sensation of how hard you are driving your leg. Look at the scale before you; we want you to use this scale from 6 to 20, where 6 means “no exertion at all” and 20 means “maximal exertion”. To help you choose a number that corresponds to how you feel within this range, consider the following. When you do not have the sensation to drive your leg, choose number 6 (“no exertion at all”). When you have the sensation to drive your leg “hard”, choose number 15. Number 20 (“Maximal exertion”) corresponds to the feeling of effort you have experienced during the preliminary maximal voluntary contraction (MVC) test. Try to appraise your perception of effort as honestly as possible, without thinking what the actual physical load is. Don’t underestimate your perception of effort but do not overestimate it either. It’s your own feeling of effort that’s important, not how it compares to other people. What other people think is not important either. Look at the scale and the expressions and then give a number. Any questions?”*

#### 2.4.2 Leg muscle pain

Leg muscle pain, defined as “the intensity of hurt that a subject feels in his quadriceps muscles only” (O’connor & Cook, 2001), was measured during the resting period between each contraction using the Cook scale (O’connor & Cook, 2001). Standardized instructions for the scale were given to each subject before the warm-up: *“You are about to undergo a resistance exercise with your knee extensor muscles (quadriceps). The scale before you contains the numbers 0 to 10. You will use this scale to assess the perceptions of pain in your quadriceps during the exercise test. For this task, pain is defined as the intensity of hurt that you feel in your quadriceps muscles only. Don’t underestimate or overestimate the degree of hurt you feel, just try to estimate it as honestly and objectively as possible. The numbers on the scale represent a range of pain intensity from “very faint pain” (number 0.5) to “extremely intense pain – almost unbearable” (number 10). When you feel no pain in your quadriceps, you should respond with the number zero. When the pain in your quadriceps becomes just noticeable, you should respond with the number 0.5. If your quadriceps feels extremely strong pain that is almost unbearable, you should respond with the number 10. You can also respond with numbers greater than 10. If the pain is greater than 10, respond with the number that represents the pain intensity you feel in relation to 10. In other words, if the pain is twice as great then respond with the number 20. Repeatedly during the test, you will be asked to rate the feelings of pain in your quadriceps. When rating these pain sensations, be sure to attend only to the specific sensations in your quadriceps and not report other pains you may be feeling (e.g., seat discomfort). It is very important that your ratings of pain intensity reflect only the degree of hurt you are feeling in your quadriceps. Do not use your ratings as an expression of fatigue (i.e., inability of the muscle to produce force) or exertion (i.e., how hard is it for you to drive your leg)*.

#### 2.4.3 Perception of force

Subjects were also asked to rate the force perceived during the contraction using a visual analogue scale ranging from 0 (“no force at all”) to 100 (“maximal force”). Participants received verbal instructions to focus specifically on the sensations of tension experienced in the muscle when providing the perception of force. Subjects were instructed that the MVC performed at the beginning of the experimental session corresponded to a force of 100. Participants were asked to report the magnitude of the force produced related to the sensations experienced during the MVC (i.e., 100 - “maximal force”).

### 2.5 Statistical analyses

To compare perceptual measurements at the same torque output, we used linear mixed-effects models (LMM; Singmann & Kellen, 2019). To test our primary deductive hypothesis that motor command is necessary for perceiving effort, we compared voluntary and evoked contractions. Additionally, to explore the relationship between the magnitude of the motor command, afferent feedback and effort perception, we compared voluntary and combined contractions. These two comparisons were estimated in separate LMMs. In each full factorial model, fixed effects included conditions (voluntary vs. evoked contractions or voluntary vs. combined contractions), group (novice vs. experienced participants), and their interaction, with by-participant random intercepts to account for the repeated-measures design. When conditions exhibited severe floor effects, i.e., near-zero variability due to predominantly zero ratings, we excluded them from the full factorial design and estimated between-group differences on the remaining conditions using LMM. Independent-samples t-tests were used to compare demographic characteristics between groups to ensure comparability. Unstandardized effect sizes were reported as β-coefficients, as they provide more context-dependent and easily interpretable results than standardized effect sizes (Singmann et al., 2023). For main effects, the β-coefficients represent half the raw mean differences between groups. When statistically significant interactions, Bonferroni-corrected post-hoc tests were conducted. Residual diagnostics for all linear mixed models were assessed using simulated residuals (Hartig & Lohse, 2022). When the residuals indicated a potential deviation from model assumptions, results were cross-validated against a robust regression approach and yielded similar estimates and conclusions. Hence, the standard models are reported throughout the manuscript. Significance level was set at α=0.05 (two-tailed) for all analyses. All statistical analyses were conducted using R version 4.3.1 (https://www.r-project.org/).

To contextualize our findings and address the constraints of our sample size (N=20), we performed simulation-based sensitivity analyses (Kumle et al., 2021; Murayama et al., 2022) and binomial probability tests. We fitted a large number of LMM models to simulated datasets (1000 simulations per effect size). Each dataset was generated with the same variance components as our observed data, but with a specific effect size (β) systematically specified. Statistical power was defined as the proportion of simulated datasets in which this effect was found to be statistically significant, with a target power of 80 ± 1%.

For the comparison of *voluntary vs. evoked contractions*, estimating the fully 2 × 2 factorial design was not feasible due to the complete absence of perceived effort reported during evoked contractions. A binomial probability analysis instead supported our finding. Under a hypothetical null hypothesis, without a priori knowledge of the true distribution of a participant’s rating of the presence of effort perception by chance, we assumed a 50% probability that a participant would report the presence of effort perception in the absence of a motor command. Considering this assumption, the probability of our true observed outcome (3—trials × 20—participants = 60 reports of no perceived effort during evoked) is extremely low (*P* = 8.67 × 10^-19^). This probability robustly confirms that the complete absence of perceived effort was not a random occurrence.

For the comparison of *voluntary vs. combined contractions*, we performed sensitivity analyses to interpret the observed effects. The primary effect of interest was the main effect of condition, i.e., the effect of manipulating the magnitude of the central motor command and the afferent feedback on the perception of effort. For this effect, given our sample size, the sensitivity analysis revealed a minimum detectable effect size of β=0.86 for the isometric contractions. For the dynamic contractions, the sensitivity analysis revealed a minimum detectable effect size of β=0.80. Importantly, the main effect of condition could be altered by the group × condition interaction, showing different trends in perceived effort between participants with or without experience with electromyostimulation (i.e., *experienced vs. novice groups)*.

As indicated in the manuscript, we also performed exploratory analyses on the group × condition interaction. For this interaction, given our sample size, the sensitivity analysis revealed a minimum detectable effect size of β=1.86 for the isometric contractions. For the dynamic contractions, the sensitivity analysis revealed a minimum detectable effect size of β=0.49.

## 3. Results

Experienced and novice participants did not differ in height (t_(18)_=0.995, p=.333), weight (t_(18)_=0.715, *p*=.484), age (t_(18)_=1.724, *p*=.102), stimulation intensity (t_(18)_=-0.192, *p*=.850) or MVC (t_(18)_=1.252, *p*=.227).

### 3.1 Present vs absent motor command

We compared voluntary contractions with evoked contractions (isometric: 10% MVC; dynamic: 5% MVC) to isolate the contributions of the motor command and afferent feedback to the generation of effort perception. Perceptions of effort, force and muscle pain are reported in **Figure 3**. During isometric contractions, the torque-time integral was similar across groups (t_(18)_=0.271, β=2.31 [-15.01, 19.61], *p*=.789) and conditions (t_(18)_=0.781, β=4.73 [-7.06, 16.50], *p*=.445), with no interactions (t_(18)_=0.319, β=1.93 [-9.86, 13.71], *p*=.753).

**Figure 3.**
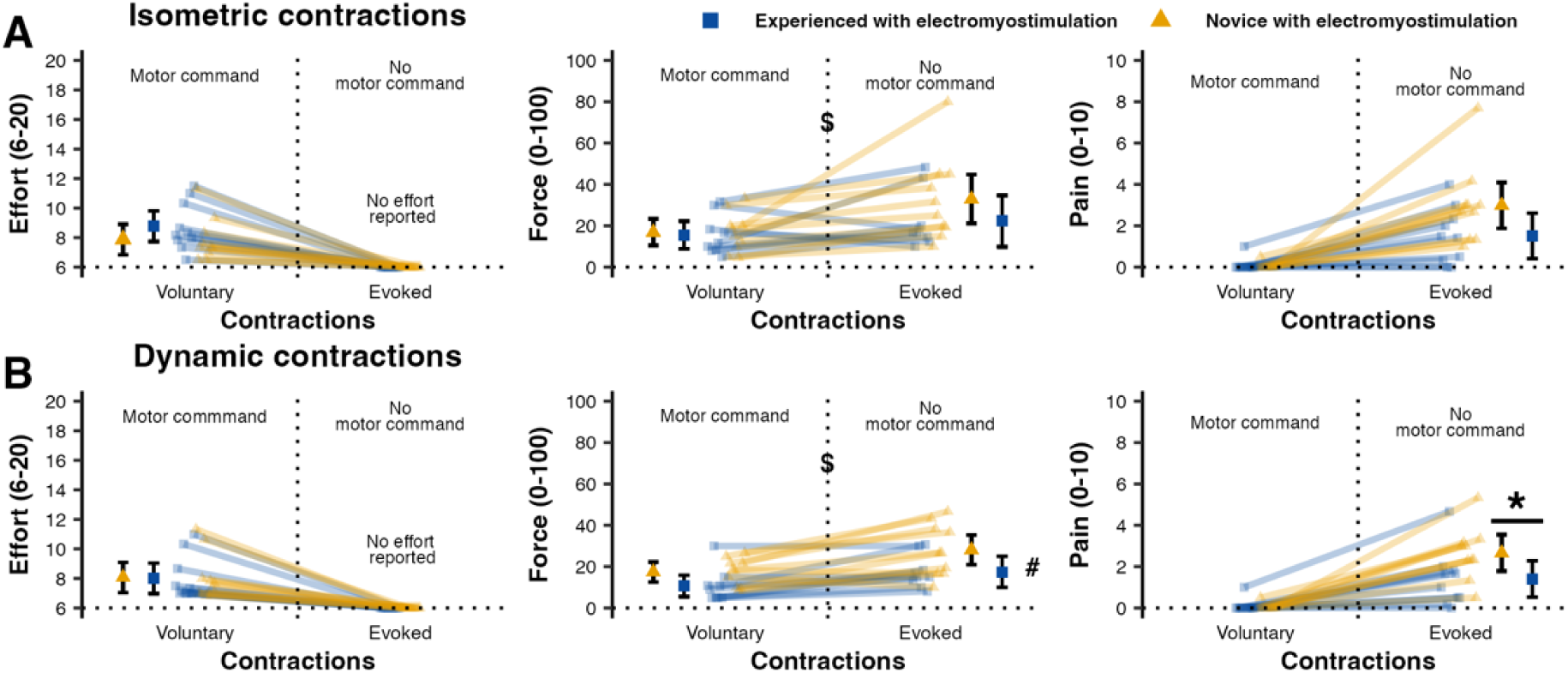
Perceptions of effort, force and muscle pain during voluntary and evoked isometric (Panel A) and dynamic (Panel B) contractions. Error bars indicate estimated marginal means ± 95% confidence intervals. Voluntary contractions involved a motor command with low afferent feedback; evoked contractions involved high afferent feedback without a motor command, matched for torque output. *N* = 20, with n = 10 per group. All plots display the full range of their respective measurement scales. $ denotes a main effect of condition; # denotes a main effect of group; * denotes a between-group interaction.

During dynamic contractions, the torque-time integral was similar across groups (t_(13)_=1.339, β=4.33 [-2.06, 10.74], *p*=.202), but higher for voluntary vs. evoked contractions (t_(17)_=3.430, β=3.25 [1.40, 5.10], *p*=.003), with no interaction (t_(17)_=1.339, β=-0.45 [-2.30, 1.39], *p*=.637).

#### 3.1.1 Perception of effort

During isometric contractions, no perceived effort was reported during evoked contractions. We therefore restricted the LMM to comparing perceived effort between groups during voluntary contractions. Perceived effort was similar between groups during voluntary contractions (t_(18)_=1.289, β=0.45 [-0.28, 1.18], *p*=.214).

During dynamic contractions, no perceived effort was reported during evoked contractions, and perceived effort was similar between groups (t_(18)_=-0.096, β=-0.03 [-0.76, 0.70], *p*=.925). For both isometric and dynamic contractions, perceived effort was reported only during voluntary contractions.

#### 3.1.2 Perception of force

During isometric contractions, experienced and novice participants reported similar force perception (main effect of group: t_(17)_=-1.100, β=-3.01 [-8.79, 2.77], *p*=.287). Participants reported lower force perception during the voluntary contractions compared to during evoked contractions (main effect of condition: t_(17)_= -3.196, β=-5.69 [-9.45, -1.93], *p*=.005). There was no group × condition interaction (t_(17)_=1.304, β=-2.32 [-1.44, 6.08], *p*=.210).

During dynamic contractions, experienced participants reported lower force perception than novices (main effect of group: t_(17)_=-2.315, β=-4.37 [-8.35, -0.39], *p*=.033). Participants reported lower force perception during the voluntary contractions compared to during evoked contractions (main effect of condition: t_(17)_=-4.370, β=-4.37 [-6.33, -2.42], *p*<.001). There was no group × condition interaction (t_(17)_=1.036, β=0.96 [-0.99, 2.91], *p*=.315).

#### 3.1.3 Muscle pain

Because most participants reported no pain during voluntary isometric and dynamic contractions, the full factorial design could not be estimated. We therefore restricted the LMM to comparing muscle pain between groups during evoked contractions. During isometric contractions, experienced participants reported lower pain intensities than novice participants during the evoked contractions, though this did not reach significance (t_(18)_=-1.993, β=-0.74 [-1.52, 0.04], *p*=.062). During dynamic contractions, experienced participants reported lower pain intensities during the evoked contractions than novice participants (t_(17)_=-2.1365, β=-0.63 [-1.25, -0.01], *p*=.047).

### 3.2 Low vs. high motor command

We compared voluntary contractions (high motor command, low afferent feedback) to combined contractions (low motor command, high afferent feedback) to investigate the relative contributions of each signal to effort perception. For both isometric and dynamic contractions, target torque was 20% MVC, with electromyostimulation accounting for 10% of the MVC during combined contractions. Perceptions of effort, force and muscle pain are reported in **Figure 4**. During isometric contractions, the torque-time integral was similar across groups (t_(18)_=0.366, β=5.35 [-23.20, 33.88], *p*=.719) and conditions (t_(18)_=0.656, β=4.99 [-9.88, 19.85], *p*=.520), with no interaction (t_(18)_=-0.319, β=2.426 [-17.30, 12.43], *p*=.754). During dynamic contractions, the torque-time integral was similar across groups (t_(17)_=1.490, β=15.48 [-4.67, 35.82], *p*=.154) and conditions (t_(17)_=1.036, β=6.540 [-5.75, 18.89], *p*=.316), with no interaction (t_(17)_=0.995, β=6.279 [-6.01, 18.63], *p*=.335).

**Figure 4.**
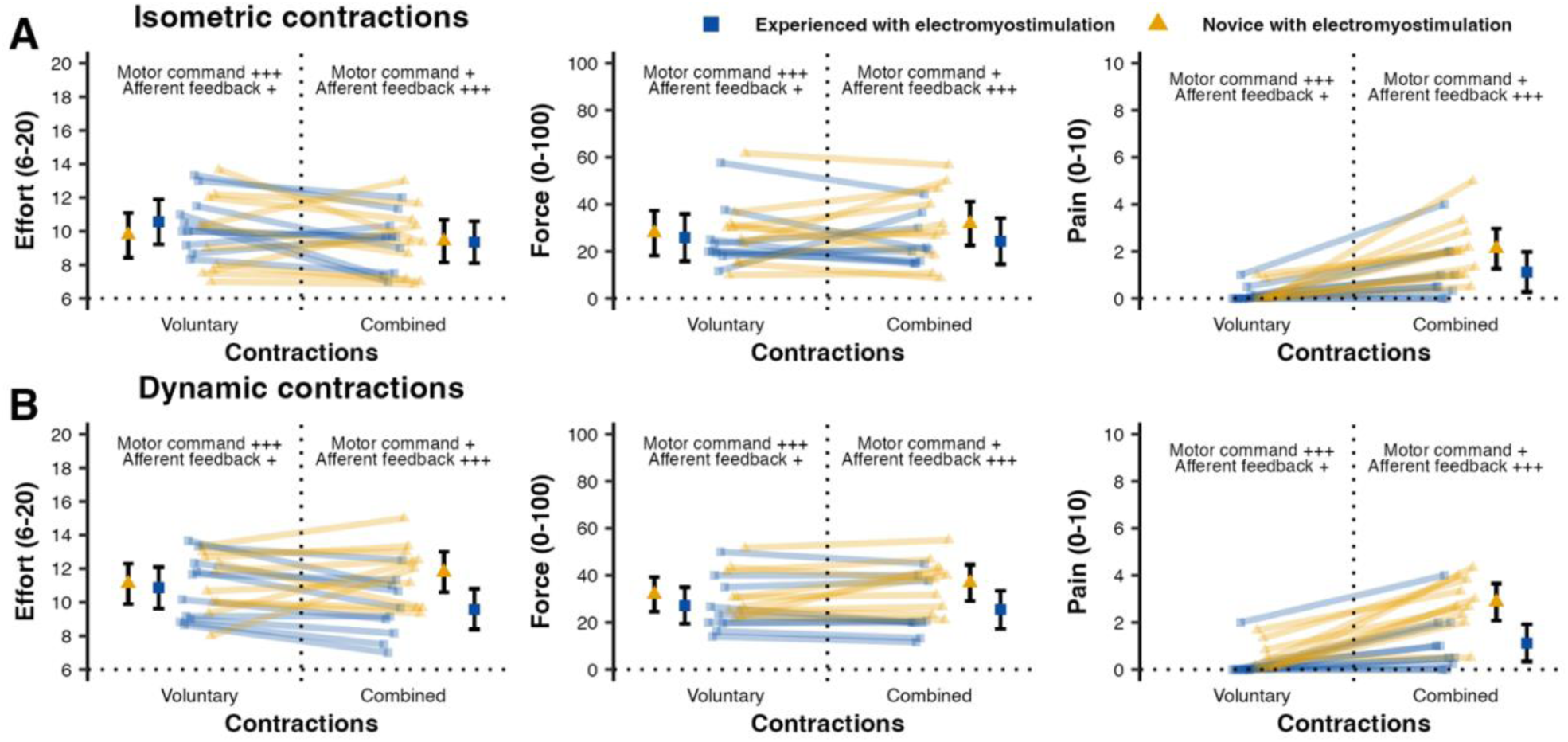
Perceptions of effort, force and muscle pain during voluntary and combined isometric (Panel A) and dynamic (Panel B) contractions. Error bars indicate estimated marginal means ± 95% confidence intervals derived from the standard error. Voluntary contractions involved high motor command with relatively low afferent feedback; combined contractions involved reduced motor command with increased afferent feedback, matched for torque output. *N* = 20, with n = 10 per group. All plots display the full range of their respective measurement scales. * denotes a between-group interaction. ▪ denotes a within-group interaction.

#### 3.2.1 Perception of effort

During isometric contractions, perceived effort was higher during voluntary contractions than combined contractions, though this did not reach the significance (main effect of condition: t_(18)_=-1.867, β=0.39 [-0.05, 0.83], *p*=.078). There was no interaction (t_(18)_=0.995, β=0.21 [-0.23, 0.65], *p*=.333).

During dynamic contractions, an interaction was observed (t_(18)_=3.012, β=0.49 [0.15, 0.83], *p*=.007). Perceived effort was similar between groups during voluntary contractions (t_(18)_=-0.301, β=-0.25 [-2.55, 2.05], *p*>.999), but experienced participants reported lower perceived effort than novices during combined contractions, although this did not reach significance (t_(18)_=-2.737, β=-2.21 [-4.44, -0.03], *p*=.054). Experienced participants reported higher perceived effort during voluntary vs. combined contractions, although this difference likewise did not reach significance (t_(18)_=2.769, β=1.27 [-0.001, 2.55], *p*=.050), while no difference was observed in novices (t_(18)_=- 1.489, β=-0.68 [-1.96, 0.59], *p*=.615).

#### 3.2.2 Perception of force

Experienced and novice participants did not report force perception differently during the isometric contractions (main effect of group: t_(17)_=-0.804, β=-2.34 [-8.50, 3.81], *p*=.433). Participants did not overall report different force perception between conditions (main effect of condition: t_(17)_=-0.434, β=-0.61 [-3.59, 2.36], *p*=.669). There was no group × condition interaction (t_(17)_=0.959, β=1.35 [-1.62, 4.33], *p*=.351).

For dynamic contractions, a group × condition interaction (t_(17)_=3.192, β=1.69 [0.64, 2.74], *p*=.002) revealed that novices reported lower force perception during purely voluntary contractions compared to combined contractions, although this did not reach significance (t_(17)_=-2.469, β=-4.90 [-11.09, 1.29], *p*=.098). No such difference was observed in experienced participants (t_(17)_=0.086, β=1.85 [-4.67, 8.38], *p*>.999).

#### 3.2.3 Muscle pain

Because most participants reported no pain during voluntary isometric and dynamic contractions, the full factorial design could not be estimated. We therefore restricted the LMM to comparing muscle pain between groups during evoked contractions. During the isometric contractions, experienced and novice participants reported similar pain intensities (t_(18)_=-1.707, β=-0.49 -1.11, 0.13], *p*=.105). During dynamic contractions, experienced participants reported lower pain intensity than novices (t_(18)_=-3.285, β=-0.87 [-1.43, -0.31], *p*=.004).

## 4. Discussion

This study aimed to test the validity of the corollary discharge and afferent feedback models in the generation of the perception of effort. To the best of our knowledge, this study is the first to use electromyostimulation to produce similar torque across contractions with different magnitudes of the central motor command and afferent feedback, and to compare ratings of perceptions of effort, force, and pain. Our data is inconsistent with the afferent feedback model. Despite stimulation of both group I-IV muscle afferents, participants reported no perception of effort in the absence of a central motor command.

### 4.1 The motor command is necessary to perceive effort

The presence of muscle pain (Pollak et al., 2014) and force (Proske & Gandevia, 2012) perception during evoked contractions without a central motor command is consistent with the successful stimulation of muscle afferents and thus, the presence of muscle feedback. We observed slightly higher torque-time integral during voluntary dynamic contractions compared to evoked contractions; a difference that was not observed during the isometric contractions. Higher torque during voluntary contractions may have occurred because participants were blind to the torque display and may have initially overshot the force required to move the dynamometer arm. Participants reported perceiving pain and force, indicating that the central nervous system processed muscle feedback during evoked contractions. Despite this sensory input and the mechanical work that was sufficient to move the arm, participants consistently rated evoked contractions as effortless. This suggests that muscle force production and muscle afferent feedback do not generate effort perception without a central motor command. The absence of perceived effort during evoked contractions aligns with a prior study showing that metabolic injections without voluntary muscle contractions can induce painful (e.g., itching, burning) and fatigue (e.g., tired, exhausted) sensations, but not effort (Pollak et al., 2014). In contrast, perceived effort is reported in the presence of a motor command without muscle feedback. For instance, participants report effort perception during imagined knee extensions, suggesting that the intention to engage in a task is a prerequisite for the experience of effort (Jacquet et al., 2021). Similarly, deafferented patients who lack muscle feedback still perceive effort (Lafargue et al., 2003). Altogether, these observations indicate that effort perception is not a passive monitoring of motor output, but a reflection of intentional engagement in a task (Mangin & Pageaux, 2026). Furthermore, experienced participants reported lower perceived effort during the combined contractions, which required a lower central motor command. Recent studies employing electrical stimulation during cycling tasks further corroborate this relationship between perceived effort and the central motor command. During cycling, increasing power output via electrical stimulation does not increase perceived effort when voluntary contraction intensity is held constant (Descollonges et al., 2025; Watanabe et al., 2021, 2023). Collectively, these results underscore the role of the central motor command in perceiving effort and challenge findings from studies reporting perceived effort at rest (Brownstein et al., 2021). Such ratings likely reflect unclear instructions or instructions that include sensations such as pain and discomfort that can be experienced even at rest (Bergevin et al., 2023).

### 4.2 Effort perception scales with the magnitude of the motor command

A notable finding is the distinct responses between experienced and novice participants in perceived effort during combined contractions, reflected in a significant group × condition interaction during dynamic contractions, in which the contribution of electromyostimulation allowed for a lower central motor command to maintain target torque than in voluntary contractions. This is broadly consistent with existent evidence that perceived effort scales with the magnitude of the central motor command (De Morree et al., 2012; Kozlowski et al., 2021). Our results suggest that experienced participants perceived a lower effort during combined compared to voluntary contractions, whereas novices did not. However, the specific comparisons did not reach significance, likely due to limited power of the secondary objective (see Limitations). Therefore, future studies adequately powered to address this question as a primary objective are needed to clarify this effect. This finding can be explained in two ways. Firstly, novice participants reported higher pain intensities following electromyostimulation. As pain is known to be costly (Torta et al., 2017), performing more painful muscle contractions requires more cognitive effort to inhibit the natural response to avoid pain. The feeling of effort associated with such inhibitory control (Mangin & Pageaux, 2026; Preston & Wegner, 2009) may have compensated for the lower motor effort during combined contractions in the novice group, resulting in no change in the overall perception of effort. Interestingly, inhibitory control is associated with the activity of premotor and motor areas (Wessel & Anderson, 2023), suggesting that the perception of effort during physical and cognitive exertion may share similar centrally generated signals. Secondly, novices may have required more cognitive control of movement to coordinate torque production during combined contractions, especially during dynamic contractions as they were not used to perform such contractions under the influence of electrostimulation. The feeling of effort associated with this potential additional cognitive control may have masked the reduction in perceived effort expected with the lower descending motor drive required to generate the combined contraction. Although we utilized large electrode (50 cm^2^) to minimize discomfort (Flodin et al., 2022) novice participants still reported higher pain intensities during dynamic contractions. Clinically, this underscores that while standard large-pad placements are pragmatic and widely used (Flodin et al., 2022), the resulting deep muscular pain may initially increase the overall perceived cost of an exercise for novices (Mangin et al., 2026). Conversely, future experimental research seeking to remove the confound of pain should map the motor point, allowing targeted muscle activation at much lower, non-painful current intensities (Gobbo et al., 2014).

### 4.3 Afferent feedback modulates central motor command but does not generate the perception of effort

Afferent feedback is known to be involved in the perception of pain (Pollak et al., 2014) and force (Proske & Gandevia, 2012), and experience during electromyostimulation confirms its stimulation. Muscle pain, primarily generated through the integration of group III/IV muscle afferent (Pollak et al., 2014), was notably minimal during voluntary contractions, but pronounced during evoked contractions. This discrepancy likely arises because the low intensities used for voluntary contractions (5% and 10% MVC) are not sufficient to elicit significant pain. Conversely, pain experienced during electromyostimulation underscores the stimulation of group III/IV muscle afferents (Bergquist et al., 2011; Pollak et al., 2014). The brain processes signals from group I/II muscle afferents to perceive force, potentially adjusting this perception based on comparisons with the central motor command (Proske & Gandevia, 2012). Our findings indicate that force perception was lower when a central motor command was present, consistent with previous research suggesting that the generation of the central motor command attenuates force perception (Voss et al., 2006). Crucially, despite the engagement of both group I-IV muscle afferents, the perception of effort is not generated without the presence of a central motor command. Neuroimaging evidence confirms that electromyostimulation activates the sensorimotor network via ascending afferent input, without any voluntary motor command (Wegrzyk et al., 2017). Despite this cortical activation, afferent feedback failed to generate effort perception. This is consistent with results from a metabolite injection study in resting muscle, which elicited pain and fatigue but not a sense of effort (Pollak et al., 2014). This does not imply that effort perception is not influenced by afferent feedback, but rather that afferent feedback indirectly contributes to effort perception via its role in motor control and in modulating the motor command (Bergevin et al., 2023, p. 416; Mangin & Pageaux, 2026, p. 407; Marchand et al., 2025; Proske & Gandevia, 2012). For example, increase in muscle spindle activity via tendon vibration, alters muscle feedback, leading to adjustment of central motor command required to execute the task, thereby modulating perceived effort (Marchand et al., 2025; McCloskey et al., 1974). Furthermore, muscle spindle reafferent feedback is involved in motor control, both at the spinal level, which mediates rapid adjustments such as the stretch reflex, and at the supraspinal level, where the actual consequences of the movement are compared with the intended motor command (Proske & Gandevia, 2012). Consistent with a spinal-level contribution, superimposing a voluntary contraction onto an evoked contraction has been shown to increase H-reflex amplitude, indicating increased spinal excitability without corresponding change of the M-wave (Borzuola et al., 2023). Facilitating muscle spindle feedback via Ia afferents increases motor pathway excitability and motor unit recruitment (Monjo et al., 2018; Monjo & Forestier, 2018). These mechanisms will alter the motor command and, therefore, the corollary discharge. Group III-IV muscle afferents contribute to the development of neuromuscular fatigue (Amann et al., 2020) and the experience of pain (O’Connor & Cook, 1999; O’Malley et al., 2024). Both phenomena can alter spinal and supraspinal mechanisms (Amann et al., 2020; Rohel et al., 2021), requiring an increase in central motor command to maintain performance. Consequently, muscle afferent feedback can indirectly contribute to the perception of effort by regulating the motor command required to execute a task.

### 4.4 Effort perception can be dissociated from other exercise-related perceptions

Finally, effort was perceived in the absence of pain during low-intensity contractions. In contrast, participants experienced pain without concurrent perception of effort during the evoked contraction. Moreover, we observed an increase in force perception during evoked contractions, compared to voluntary contractions, without a corresponding increase in reported effort. These findings are consistent with the existing literature, which supports the idea that individuals can dissociate perceptions of effort from other exercise-related perceptions (Pageaux, 2016). These discrete experiences emphasize the complexity of sensory processing during engagement in physical tasks and reinforce the idea that the perception of effort is a distinct phenomenon (see Bergevin et al., 2023; Halperin & Emanuel, 2020; Pageaux, 2016).

### 4.5 Limitations and perspectives

Our study was primarily designed to test the hypothesis that the generation of effort perception strictly depends on the presence of a central motor command. Accordingly, the sample size was determined for this purpose (Abt et al., 2025; Ditroilo et al., 2025), and our results suggest that effort is not generated in the absence of volitional drive. Our secondary objective, examining the relationship between the magnitude of the central motor command and effort perception, was not the basis for this sample size. Statistical power for this secondary comparison was therefore likely lower, particularly for the post-hoc comparisons of the significant of the group × condition interactions. In this context, we encourage future studies to replicate our experimental manipulation of motor command magnitude and afferent feedback during voluntary and combined contractions with larger sample sizes. A second limitation is the lack of a neurophysiological measure to quantify the magnitude of the central motor command and muscle afferent feedback. While our behavioural results align with the corollary discharge model, we did not record the movement-related cortical potential (De Morree et al., 2012; Shibasaki & Hallett, 2006) or other cortical markers that would provide an objective measure of volitional drive. Similarly, we did not use microneurography to measure the magnitude of muscle afferent feedback (Kakuda & Nagaoka, 1998). We are nonetheless confident in the interpretation of our exploratory analyses regarding the relationship between effort perception and the magnitude of the central motor command. Previous psychophysiological research supports the view that an increase in muscle contraction intensity requires a concomitantly stronger motor command (Berchicci et al., 2013; De Morree et al., 2012). During the combined contractions, participants had to produce only 10% of their MVC force (the other 10% was produced with electromyostimulation), compared to 20% during voluntary contractions. According to Henneman’s size principle (Pageaux & Lepers, 2016), these submaximal intensities primarily recruit low-threshold motor units. However, doubling the voluntary torque output still necessitates a distinct, measurable increase in the magnitude of the central motor command. We deliberately selected these intensities to ensure that ratings of perceived effort were not confounded by fatigue (Pageaux & Lepers, 2016) or high pain levels. Further investigation should incorporate electroencephalography-based measures, such as motor-related cortical potentials, or compare concentric and eccentric contractions to manipulate the magnitude of the motor command while minimizing peripheral confounding factors.

Our results, combined with others (De Morree et al., 2012; Kozlowski et al., 2021), suggest that central motor command (defined as activity of premotor and motor cortical areas) is essential for the generation of perceived effort, whilst afferent feedback is not. The results in novices who did not report lower ratings of perceived effort during combined contractions suggest that the activity of premotor and motor areas probably related to inhibitory control and cognitive control of movement, and not just the magnitude of the descending motor drive required for the muscle contraction, contribute to the perception of effort experienced during physical tasks. This view is supported by studies showing that the cortical areas associated with the perception of effort during physical tasks such as the supplementary motor area (Zénon et al., 2015) and the anterior cingulate cortex (Williamson et al., 2001) are upstream of the primary motor cortex and also associated with cognitive control (Gillies et al., 2019; Nachev et al., 2008; Shackman et al., 2011). Further exploration of the cortical circuits associated with the perceived effort during physical and cognitive exertion is required. Understanding these neural underpinnings is crucial, particularly because the deployment of effort is intrinsically a choice that can affect human behaviour and sporting performance, among other things (Marcora & Staiano, 2010).

### 4.6 Conclusion

To conclude, our results reinforce the view that, to generate the perception of effort, the brain primarily processes signals related to the motor command rather than afferent feedback. Even if afferent feedback does not seem to be the sensory signal processed by the brain to produce the conscious experience of effort, it can influence the perception of effort indirectly. For example, by generating the conscious experience of pain requiring effortful inhibitory control, or by inhibiting the motor neurons, thus forcing an increase in central command to produce the same submaximal force. A clearer understanding of the neural mechanisms underlying effort perception is crucial for developing targeted interventions to improve engagement and adherence to prescribed activity (e.g., physical exercise; Marcora, 2016). For instance, effort perception is heightened in several conditions, such as post-stroke patients (Kuppuswamy et al., 2015), cancer patients (Fernandez et al., 2020) or kidney patients (Macdonald et al., 2012), acting as a substantial barrier to rehabilitation. Recent evidence suggests that reducing the required motor command, such as in a cycling task, can lower perceived effort in clinical populations (Descollonges et al., 2025). Ultimately, mapping the neural pathway of effort perception can better inform clinical practice and improve patient care.

## Ethics approval

All experimental protocols and procedures were approved by the local ethics committee. All participants provided informed consent prior to the study.

## Author Contributions

BP, RL and SM designed the study. BP and LA collected the data. MB and TM performed the statistical analyses. MB and BP created the figures. MB and TM wrote the first draft of the manuscript. All authors revised the first draft and approved the final version of the manuscript.

## Use of AI-Generated Tools

The authors used Gemini 3.1 Pro and ChatGPT 5.2 to assist with language editing during manuscript preparation. The authors reviewed and edited the manuscript content as needed and take full responsibility for the content of the published article. All scientific content, data interpretation, and conclusions were generated solely by the authors.

## References

Abt, G., Boreham, C., Davison, G., Jackson, R., Jobson, S., Wallace, E., & Williams, M. (2025). Sample size estimation revisited. Journal of Sports Sciences, 1–6. DOI.org (Crossref). 10.1080/02640414.2025.2499403

Alon, G., & Smith, G. (2005). Tolerance and conditioning to neuro-muscular electrical stimulation within and between sessions and gender. Journal of Sports Science & Medicine, 4(4), 395–405.

Amann, M., Blain, G. M., Proctor, L. T., Sebranek, J. J., Pegelow, D. F., & Dempsey, J. A. (2010). Group III and IV muscle afferents contribute to ventilatory and cardiovascular response to rhythmic exercise in humans. Journal of Applied Physiology, 109(4), 966–976. DOI.org (Crossref). 10.1152/japplphysiol.00462.2010

Amann, M., Wan, H.-Y., Thurston, T. S., Georgescu, V. P., & Weavil, J. C. (2020). On the Influence of Group III/IV Muscle Afferent Feedback on Endurance Exercise Performance. Exercise and Sport Sciences Reviews, 48(4), 209. journals.lww.com. 10.1249/JES.0000000000000233

Berchicci, M., Menotti, F., Macaluso, A., & Di Russo, F. (2013). The neurophysiology of central and peripheral fatigue during sub-maximal lower limb isometric contractions. Frontiers in Human Neuroscience, 7. DOI.org (Crossref). 10.3389/fnhum.2013.00135

Bergevin, M., Steele, J., Payen de la Garanderie, M., Feral-Basin, C., Marcora, S. M., Rainville, P., Caron, J. G., & Pageaux, B. (2023). Pharmacological Blockade of Muscle Afferents and Perception of Effort: A Systematic Review with Meta-analysis. Sports Medicine, 53(2), 415–435. Springer Link. 10.1007/s40279-022-01762-4

Bergquist, A. J., Clair, J. M., Lagerquist, O., Mang, C. S., Okuma, Y., & Collins, D. F. (2011). Neuromuscular electrical stimulation: Implications of the electrically evoked sensory volley. European Journal of Applied Physiology, 111(10), 2409–2426. DOI.org (Crossref). 10.1007/s00421-011-2087-9

Bingel, U., Schoell, E., Herken, W., Büchel, C., & May, A. (2007). Habituation to painful stimulation involves the antinociceptive system. Pain, 131(1), 21–30. DOI.org (Crossref). 10.1016/j.pain.2006.12.005

Boerio, D., Jubeau, M., Zory, R., & Maffiuletti, N. A. (2005). Central and Peripheral Fatigue after Electrostimulation-Induced Resistance Exercise. Medicine & Science in Sports & Exercise, 37(6), 973. journals.lww.com. 10.1249/01.mss.0000166579.81052.9c

Borg, G. (1998). Borg’s perceived exertion and pain scales. Human kinetics.

Borzuola, R., Quinzi, F., Scalia, M., Pitzalis, S., Di Russo, F., & Macaluso, A. (2023). Acute effects of neuromuscular electrical stimulation on cortical dynamics and reflex activation. Journal of Neurophysiology, 129(6), 1310–1321. 10.1152/jn.00085.2023

Briand, J., Mangin, T., Tremblay, J., & Pageaux, B. (2025). Bridging Inductive and Deductive Reasoning: A Proposal to Enhance the Evaluation and Development of Models in Sports and Exercise Science. Sports Medicine, 55(11), 2707–2719. (WOS:001541709500001). 10.1007/s40279-025-02289-0

Brinkmann, K., & Gendolla, G. H. E. (2008). Does depression interfere with effort mobilization? Effects of dysphoria and task difficulty on cardiovascular response. Journal of Personality and Social Psychology, 94(1), 146–157. DOI.org (Crossref). 10.1037/0022-3514.94.1.146

Brownstein, C. G., Rimaud, D., Singh, B., Fruleux-Santos, L.-A., Sorg, M., Micklewright, D., & Millet, G. Y. (2021). French Translation and Validation of the Rating-of-Fatigue Scale. Sports Medicine - Open, 7(1), 25. BioMed Central. 10.1186/s40798-021-00316-8

Broxterman, R. M., Hureau, T. J., Layec, G., Morgan, D. E., Bledsoe, A. D., Jessop, J. E., Amann, M., & Richardson, R. S. (2018). Influence of group III/IV muscle afferents on small muscle mass exercise performance: A bioenergetics perspective: Influence of muscle afferents on exercise performance. The Journal of Physiology, 596(12), 2301–2314. DOI.org (Crossref). 10.1113/JP275817

Craig, A. D. (2003). Interoception: The sense of the physiological condition of the body. Current Opinion in Neurobiology, 13(4), 500–505.

De Morree, H. M., Klein, C., & Marcora, S. M. (2012). Perception of effort reflects central motor command during movement execution: Neurophysiology of perceived effort. Psychophysiology, 49(9), 1242–1253. DOI.org (Crossref). 10.1111/j.1469-8986.2012.01399.x

De Morree, H. M., Klein, C., & Marcora, S. M. (2014). Cortical substrates of the effects of caffeine and time-on-task on perception of effort. Journal of Applied Physiology, 117(12), 1514–1523. DOI.org (Crossref). 10.1152/japplphysiol.00898.2013

Descollonges, M., Di Marco, J., Jafari, E., Pouillart, P.-H., Brugniaux, J. V., Pageaux, B., & Deley, G. (2025). Positive effects of functional electrical stimulation-assisted cycling on perception of effort, cerebral blood flow, and cognition in post-stroke patients. Journal of NeuroEngineering and Rehabilitation, 22(1), 257. DOI.org (Crossref). 10.1186/s12984-025-01800-y

Dimitriou, M. (2022). Human muscle spindles are wired to function as controllable signal-processing devices. eLife, 11, e78091. 10.7554/eLife.78091

Ditroilo, M., Mesquida, C., Abt, G., & Lakens, D. (2025). Exploratory research in sport and exercise science: Perceptions, challenges, and recommendations. Journal of Sports Sciences, 1–13. PubMed. 10.1080/02640414.2025.2486871

Fernandez, C., Firdous, S., Jehangir, W., Behm, B., Mehta, Z., Berger, A., & Davis, M. (2020). Cancer-Related Fatigue: Perception of Effort or Task Failure? American Journal of Hospice and Palliative Medicine, 37(1), 34–40. 10.1177/1049909119849420

Flodin, J., Juthberg, R., & Ackermann, P. W. (2022). Effects of electrode size and placement on comfort and efficiency during low-intensity neuromuscular electrical stimulation of quadriceps, hamstrings and gluteal muscles. BMC Sports Science, Medicine and Rehabilitation, 14(1), 11. 10.1186/s13102-022-00403-7

Gillies, M. J., Huang, Y., Hyam, J. A., Aziz, T. Z., & Green, A. L. (2019). Direct neurophysiological evidence for a role of the human anterior cingulate cortex in central command. Autonomic Neuroscience, 216, 51–58. DOI.org (Crossref). 10.1016/j.autneu.2018.09.004

Gobbo, M., Maffiuletti, N. A., Orizio, C., & Minetto, M. A. (2014). Muscle motor point identification is essential for optimizing neuromuscular electrical stimulation use. Journal of NeuroEngineering and Rehabilitation, 11(1), 17. 10.1186/1743-0003-11-17

Gruet, M., Mely, L., & Vallier, J. M. (2018). Overall and differentiated sensory responses to cardiopulmonary exercise test in patients with cystic fibrosis: Kinetics and ability to predict peak oxygen uptake. Eur J Appl Physiol, 118(9), 2007–2019. Nlm. 10.1007/s00421-018-3923-y

Grummt, M., Hafermann, L., Claussen, L., Herrmann, C., & Wolfarth, B. (2024). Rating of Perceived Exertion: A Large Cross-Sectional Study Defining Intensity Levels for Individual Physical Activity Recommendations. Sports Medicine - Open, 10(1), 71. DOI.org (Crossref). 10.1186/s40798-024-00729-1

Halperin, I., & Emanuel, A. (2020). Rating of Perceived Effort: Methodological Concerns and Future Directions. Sports Medicine, 50(4), 679–687. Springer Link. 10.1007/s40279-019-01229-z

Hartig, F., & Lohse, L. (2022). DHARMa: Residual Diagnostics for Hierarchical (Multi-Level / Mixed) Regression Models [Computer software]. R-Packages.

Jacquet, T., Lepers, R., Poulin-Charronnat, B., Bard, P., Pfister, P., & Pageaux, B. (2021). Mental fatigue induced by prolonged motor imagery increases perception of effort and the activity of motor areas. Neuropsychologia, 150, 107701. DOI.org (Crossref). 10.1016/j.neuropsychologia.2020.107701

Kakuda, N., & Nagaoka, M. (1998). Dynamic response of human muscle spindle afferents to stretch during voluntary contraction. The Journal of Physiology, 513(2), 621–628. DOI.org (Crossref). 10.1111/j.1469-7793.1998.621bb.x

Kozlowski, B., Pageaux, B., Hubbard, E. F., St. Peters, B., Millar, P. J., & Power, G. A. (2021). Perception of effort during an isometric contraction is influenced by prior muscle lengthening or shortening. European Journal of Applied Physiology, 121(9), 2531–2542. DOI.org (Crossref). 10.1007/s00421-021-04728-y

Kumle, L., Võ, M. L. H., & Draschkow, D. (2021). Estimating power in (generalized) linear mixed models: An open introduction and tutorial in R. Behavior Research Methods, 53(6), 2528–2543. Springer Link. 10.3758/s13428-021-01546-0

Kuppuswamy, A., Clark, E. V., Turner, I. F., Rothwell, J. C., & Ward, N. S. (2015). Post-stroke fatigue: A deficit in corticomotor excitability? Brain, 138(1), 136–148. 10.1093/brain/awu306

Lafargue, G., Paillard, J., Lamarre, Y., & Sirigu, A. (2003). Production and perception of grip force without proprioception: Is there a sense of effort in deafferented subjects?: Sensation of effort after deafferentation. European Journal of Neuroscience, 17(12), 2741–2749. DOI.org (Crossref). 10.1046/j.1460-9568.2003.02700.x

Lakens, D. (2022). Sample Size Justification. Collabra: Psychology, 8(1), 33267. DOI.org (Crossref). 10.1525/collabra.33267

Macdonald, J. H., Fearn, L., Jibani, M., & Marcora, S. M. (2012). Exertional Fatigue in Patients With CKD. American Journal of Kidney Diseases, 60(6), 930–939. 10.1053/j.ajkd.2012.06.021

Mangin, T., Monti, I., Marcotte, M., Baudry, S., Roy, M., Rainville, P., & Pageaux, B. (2026). Maintaining performance under pain is effortful: Experimental and computational evidence. bioRxiv, 2026.02.13.705857. 10.64898/2026.02.13.705857

Mangin, T., & Pageaux, B. (2026). Effort and its perception revisited: How physical-domain insights could lead toward a unified theory. Cognitive, Affective, & Behavioral Neuroscience. 10.3758/s13415-026-01411-7

Marchand, F., Pageaux, B., Forestier, N., & Monjo, F. (2025). Prolonged passive vibration of Achilles and patellar tendons decreases effort perception during subsequent cycling tasks. Journal of Sport and Health Science, 101061. DOI.org (Crossref). 10.1016/j.jshs.2025.101061

Marcora, S. M. (2009). Perception of effort during exercise is independent of afferent feedback from skeletal muscles, heart, and lungs. Journal of Applied Physiology, 106(6), 2060–2062. 10.1152/japplphysiol.90378.2008

Marcora, S. M. (2010). Perception: Effort of. In Encyclopedia of perception (Goldstein, E. B., pp. 380–383). Sage.

Marcora, S. M. (2016). Can Doping be a Good Thing? Using Psychoactive Drugs to Facilitate Physical Activity Behaviour. Sports Med, 46(1), 1–5. Nlm. 10.1007/s40279-015-0412-x

Marcora, S. M., & Staiano, W. (2010). The limit to exercise tolerance in humans: Mind over muscle? European Journal of Applied Physiology, 109(4), 763–770. 10.1007/s00421-010-1418-6

Mater, A., Boly, A., Assadi, H., Martin, A., & Lepers, R. (2023). Effect of Cadence on Physiological and Perceptual Responses during Eccentric Cycling at Different Power Outputs. Medicine & Science in Sports & Exercise, 55(6), 1105–1113. DOI.org (Crossref). 10.1249/MSS.0000000000003132

McCloskey, D. I., Ebeling, P. R., & Goodwin, G. M. (1974). Estimation of weights and tensions and apparent involvement of a “sense of effort.” Experimental Neurology, 42(1), 220–232. 10.1016/0014-4886(74)90019-3

Monjo, F., & Allen, T. (2023). What if muscle spindles were also involved in the sense of effort? The Journal of Physiology, JP284376. DOI.org (Crossref). 10.1113/JP284376

Monjo, F., & Forestier, N. (2018). Muscle spindle thixotropy affects force perception through afferent-induced facilitation of the motor pathways as revealed by the Kohnstamm effect. Experimental Brain Research, 236(4), 1193–1204. Springer Link. 10.1007/s00221-018-5207-5

Monjo, F., Shemmell, J., & Forestier, N. (2018). The sensory origin of the sense of effort is context-dependent. Experimental Brain Research, 236(7), 1997–2008. Springer Link. 10.1007/s00221-018-5280-9

Murayama, K., Usami, S., & Sakaki, M. (2022). Summary-statistics-based power analysis: A new and practical method to determine sample size for mixed-effects modeling. Psychological Methods. DOI.org (Crossref). 10.1037/met0000330

Nachev, P., Kennard, C., & Husain, M. (2008). Functional role of the supplementary and pre-supplementary motor areas. Nature Reviews Neuroscience, 9(11), 856–869. DOI.org (Crossref). 10.1038/nrn2478

Norbury, R., Smith, S. A., Burnley, M., Judge, M., & Mauger, A. R. (2022a). The effect of elevated muscle pain on neuromuscular fatigue during exercise. Eur J Appl Physiol, 122(1), 113–126. (34586471). 10.1007/s00421-021-04814-1

Norbury, R., Smith, S. A., Burnley, M., Judge, M., & Mauger, A. R. (2022b). The effect of hypertonic saline evoked muscle pain on neurophysiological changes and exercise performance in the contralateral limb. Exp Brain Res, 240(5), 1423–1434. Nlm. 10.1007/s00221-022-06342-6

O’Connor, P. J., & Cook, D. B. (1999). Exercise and Pain: The Neurobiology, Measurement, and Laboratory Study of Pain in Relation to Exercise in Humans. Exercise and Sport Sciences Reviews, 27(1), 119–166.

O’connor, P. J., & Cook, D. B. (2001). Moderate-intensity muscle pain can be produced and sustained during cycle ergometry. Medicine & Science in Sports & Exercise, 33(6), 1046. journals.lww.com.

O’Malley, C., Smith, S., Mauger, A., & Norbury, R. (2024). Exercise-induced pain within endurance exercise settings: Definitions, measurement, mechanisms and potential interventions. Experimental Physiology, 109(9), 1446–1460. DOI.org (Crossref). 10.1113/EP091687

Pageaux, B. (2016). Perception of effort in exercise science: Definition, measurement and perspectives. European Journal of Sport Science, 16(8), 885–894. 10.1080/17461391.2016.1188992

Pageaux, B., & Lepers, R. (2016). Fatigue Induced by Physical and Mental Exertion Increases Perception of Effort and Impairs Subsequent Endurance Performance. Frontiers in Physiology, 7. DOI.org (Crossref). 10.3389/fphys.2016.00587

Pollak, K. A., Swenson, J. D., Vanhaitsma, T. A., Hughen, R. W., Jo, D., Light, K. C., Schweinhardt, P., Amann, M., & Light, A. R. (2014). Exogenously applied muscle metabolites synergistically evoke sensations of muscle fatigue and pain in human subjects: Synergistic metabolites evoke muscle pain and fatigue. Experimental Physiology, 99(2), 368–380. DOI.org (Crossref). 10.1113/expphysiol.2013.075812

Preston, J., & Wegner, D. M. (2009). Elbow grease: When action feels like work. Social Cognition and Social Neuroscience. Oxford Handbook of Human Action, 569–586.

Proske, U., & Gandevia, S. C. (2012). The Proprioceptive Senses: Their Roles in Signaling Body Shape, Body Position and Movement, and Muscle Force. Physiological Reviews, 92(4), 1651–1697. DOI.org (Crossref). 10.1152/physrev.00048.2011

Rohel, A., Bouffard, J., Patricio, P., Mavromatis, N., Billot, M., Roy, J. S., Bouyer, L., Mercier, C., & Masse-Alarie, H. (2021). The effect of experimental pain on the excitability of the corticospinal tract in humans: A systematic review and meta-analysis. Eur J Pain, 25(6), 1209–1226. Nlm. 10.1002/ejp.1746

Shackman, A. J., Salomons, T. V., Slagter, H. A., Fox, A. S., Winter, J. J., & Davidson, R. J. (2011). The integration of negative affect, pain and cognitive control in the cingulate cortex. Nature Reviews Neuroscience, 12(3), 154–167. DOI.org (Crossref). 10.1038/nrn2994

Shenhav, A., Musslick, S., Lieder, F., Kool, W., Griffiths, T. L., Cohen, J. D., & Botvinick, M. M. (2017). Toward a Rational and Mechanistic Account of Mental Effort. Annual Review of Neuroscience, 40(1), 99–124. DOI.org (Crossref). 10.1146/annurev-neuro-072116-031526

Shibasaki, H., & Hallett, M. (2006). What is the Bereitschaftspotential? Clinical Neurophysiology, 117(11), 2341–2356. DOI.org (Crossref). 10.1016/j.clinph.2006.04.025

Singmann, H., & Kellen, D. (2019). An introduction to mixed models for experimental psychology. In New methods in cognitive psychology (pp. 4–31). Routledge.

Singmann, H., Kellen, D., Cox, G. E., Chandramouli, S. H., Davis-Stober, C. P., Dunn, J. C., Gronau, Q. F., Kalish, M. L., McMullin, S. D., Navarro, D. J., & Shiffrin, R. M. (2023). Statistics in the Service of Science: Don’t Let the Tail Wag the Dog. Computational Brain & Behavior, 6(1), 64–83. Springer Link. 10.1007/s42113-022-00129-2

Torta, D. M., Legrain, V., Mouraux, A., & Valentini, E. (2017). Attention to pain! A neurocognitive perspective on attentional modulation of pain in neuroimaging studies. Cortex, 89, 120–134. DOI.org (Crossref). 10.1016/j.cortex.2017.01.010

Voss, M., Ingram, J. N., Haggard, P., & Wolpert, D. M. (2006). Sensorimotor attenuation by central motor command signals in the absence of movement. Nature Neuroscience, 9(1), 26–27. www.nature.com. 10.1038/nn1592

Watanabe, K., Takada, T., Kawade, S., & Moritani, T. (2021). Effect of exercise intensity on metabolic responses on combined application of electrical stimulation and voluntary exercise. Physiological Reports, 9(3). DOI.org (Crossref). 10.14814/phy2.14758

Watanabe, K., Yoshimura, A., Nojima, H., Hirono, T., Kunugi, S., Takada, T., Kawade, S., & Moritani, T. (2023). Physiological adaptations following vigorous exercise and moderate exercise with superimposed electrical stimulation. European Journal of Applied Physiology, 123(1), 159–168. DOI.org (Crossref). 10.1007/s00421-022-05065-4

Wegrzyk, J., Ranjeva, J.-P., Fouré, A., Kavounoudias, A., Vilmen, C., Mattei, J.-P., Guye, M., Maffiuletti, N. A., Place, N., Bendahan, D., & Gondin, J. (2017). Specific brain activation patterns associated with two neuromuscular electrical stimulation protocols. Scientific Reports, 7(1), 2742. 10.1038/s41598-017-03188-9

Wessel, J. R., & Anderson, M. C. (2023). Neural mechanisms of domain-general inhibitory control. Trends Cogn Sci. Publisher (39492255). 10.1016/j.tics.2023.09.008

Williamson, J. W., McColl, R., Mathews, D., Mitchell, J. H., Raven, P. B., & Morgan, W. P. (2001). Hypnotic manipulation of effort sense during dynamic exercise: Cardiovascular responses and brain activation. Journal of Applied Physiology, 90(4), 1392–1399. DOI.org (Crossref). 10.1152/jappl.2001.90.4.1392

Zénon, A., Sidibé, M., & Olivier, E. (2015). Disrupting the supplementary motor area makes physical effort appear less effortful. Journal of Neuroscience, 35(23), 8737–8744.

